# Brainstem pathology in human narcolepsy: Neurodegeneration in the locus coeruleus in narcoleptic humans, but not in genetically narcoleptic mice or dogs

**DOI:** 10.1101/2025.04.12.648456

**Authors:** Thomas C. Thannickal, Ming-Fung Wu, Marcia E. Cornford, Jerome M. Siegel

## Abstract

Our earlier study led to the conclusion that human narcolepsy with cataplexy was caused by the loss of forebrain hypocretin (orexin) neurons in the hypothalamus. We now report that humans having narcolepsy with cataplexy also have a 46% decrease in the number and an 18% increase in the size of brainstem neuromelanin-pigmented locus coeruleus (LC) neurons and increased microglial activity in LC. However, no such changes are observed in the LC of hypocretin peptide depleted narcoleptic mice, hypocretin neuron depleted orexin-tTA/TetO-DTA (orexin-DTA) narcoleptic mice or in narcoleptic dogs. Sodium oxybate, an effective treatment for narcolepsy, decreased the size of LC norepinephrine neurons and increased the number and size of LC microglial cells in mice. Our results indicate that the autoimmune process believed to cause human narcolepsy affects both forebrain Hcrt neurons and brainstem LC norepinephrine cells. Addressing both forebrain and brainstem pathologies may improve understanding of, and treatments for, human narcolepsy.

## Main

A study of the function of hypocretin (Hcrt/orexin), a forebrain hypothalamic peptide, utilizing genetic manipulation in mice to prevent synthesis of this peptide, noted abrupt losses of muscle tone in waking, resembling the loss of muscle tone in REM sleep and in human cataplexy^1^. This led to anatomical studies of the hypothalamic forebrain of human narcoleptics, revealing a 90% loss of neurons containing Hcrt^2–4^, and to treatment of narcolepsy by administering Hcrt^5, 6^.

Cataplexy, the emotionally triggered loss of muscle tone that occurs during waking in narcoleptics, has been likened to the loss of muscle tone that accompanies REM sleep. But REM sleep and REM sleep linked muscle tone suppression has been shown to be generated by the brainstem, not by the forebrain^7^. We have shown that cessation of activity of norepinephrine neurons in the brainstem locus coeruleus (LC) is tightly linked to the suppression of muscle tone in both REM sleep and cataplexy, in narcoleptic dogs^8^. Muscle tone suppression in REM sleep does not require the hypothalamus or any other forebrain region^7, 9^, raising the question of how the loss of Hcrt neurons in the forebrain of humans is related to this unique symptom of cataplexy. We have also found “cataplexy-on” neurons found in the medulla of narcoleptic dogs^10^.

In the current study, we conducted the first histological analysis of the LC of human narcoleptics. We analyzed the LC of 11 narcolepsy with cataplexy (NT-1) patients and 5 control subjects (Extended Data Table. 1) with hematoxylin-eosin (H&E) staining to assess the number and size of neuromelanin (NM) pigmented norepinephrine containing neurons (Fig. 1a-f). We find that all human narcoleptics have brainstem pathology in LC, in addition to the previously identified loss of forebrain hypocretin. We observed a substantial reduction in the number of LC neurons with dark pigmentation in NT-1 patients (Fig.1a, far left vs. middle left), with 46% fewer NM pigmented neurons compared to control subjects (Fig. 1a, middle right vs far right (t = 8.52, df = 14 and P = 0.001), with a loss range of 28 - 68% (Fig. 1a, b, c). Furthermore, the remaining noradrenergic LC neurons in NT-1 patients were 18% larger than controls (Fig. 1c, t = 7.32, df = 14, and P = 0.01). Fig. 1d shows the distribution of NM cells in the anterior-to-posterior dimension of the LC of a NT-1 patient (90 yrs, M) and a control subject (82 yrs, M) (Fig. 1 e & f). Cell loss was most prominent in the anterior part of the LC (Fig. 1e). There was no significant correlation between the percentage loss of NM pigmented neurons and cataplexy onset age (r = 0.28, P = 0.39).

**Fig. 1:**
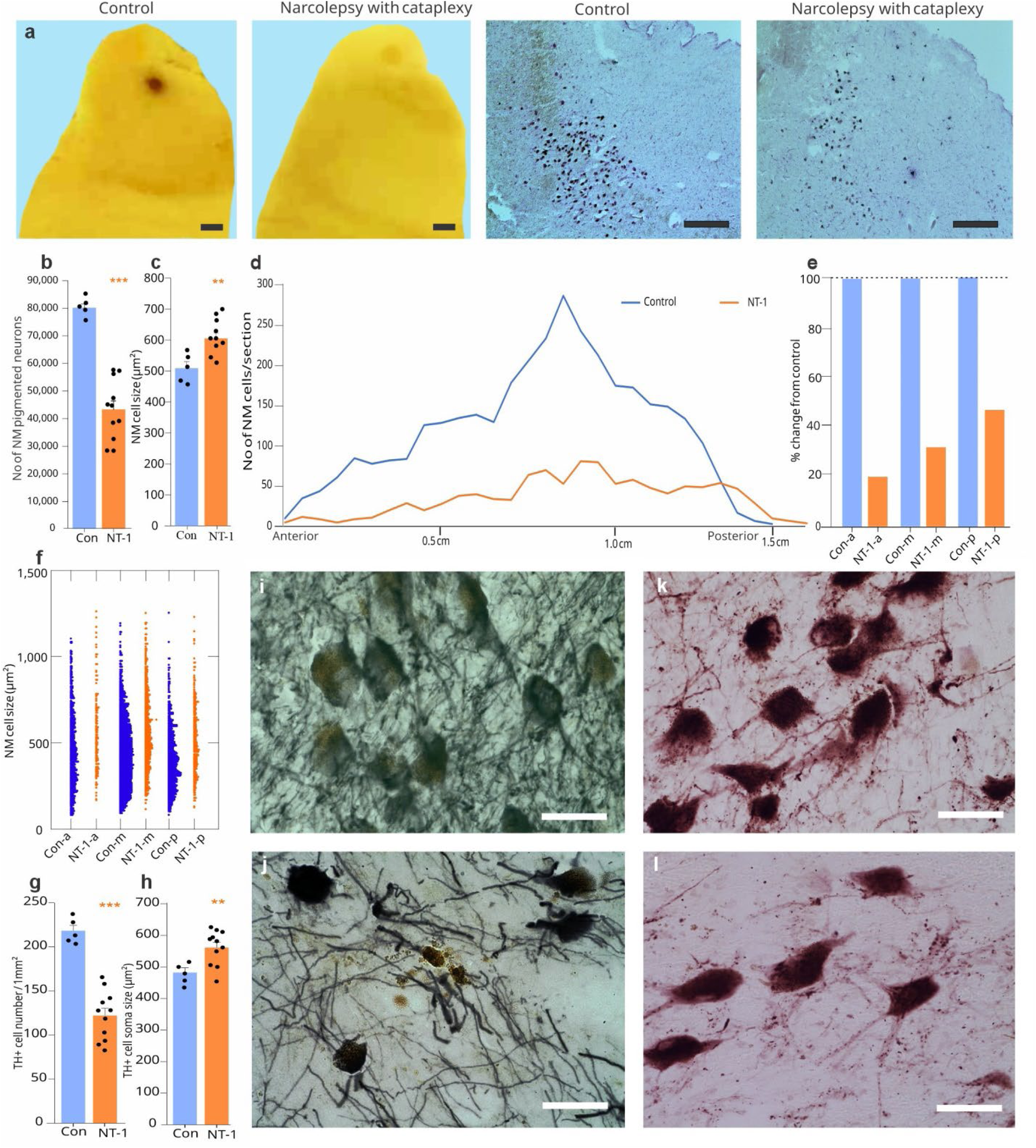
Neuromelanin and norepinephrine cell loss in the locus coeruleus of narcolepsy with cataplexy patients (NT-1). **a**, Far left, unstained brain stem block of control patient shows the dark spot of the locus coeruleus neurons, 2 – 2.5 mm in diameter. Left, center, similar unstained block shows faded locus coeruleus spot in patient of narcolepsy with cataplexy. Right, center crystal violet-stained microscopic image from control patient shows neuromelanin containing locus coeruleus neurons. Far right, narcolepsy with cataplexy patients show a reduced number of neuromelanin (NM) neurons in the locus coeruleus compared to control subjects. Fourth ventricle is to the right in each of the top 4 images. **b**, Stereological calculations show a significant reduction in the number of NM pigmented cells in narcolepsy with cataplexy patients (NT-1) (n = 11) compared to controls (n = 5) (t = 7.32, P = 0.01). **c**, Neuromelanin cell size was significantly increased in NT-1 patients (t = 3.24, P = 0.001). **d**, Neuromelanin cell distribution from anterior to posterior part of locus coeruleus of control and NT-1 patients. **e**, The percentage changes in number within each subregion in narcoleptic compared to control: (a) anterior, (m) middle, and (p) posterior in control (blue) and NT-1 (orange) subjects. **f,** NM cell size distribution from anterior, middle and posterior parts of the LC in control (Con-) and narcoleptic (NT-1) subjects. **g**, In NT-1 patients, TH+ cell density is significantly reduced compared to control (n = 11 and 5 respectively, t = 7.32, P = 0.001). **h**, Size of the TH+ neurons are larger in NT-1 than in controls (t = 2.91. P = 0.01). **i** & **j**, Images of NM pigmented neurons in control (Top) and NT-1 (bottom) with H&E counterstaining. **k** & **l**, Immunohistochemistry images of dopamine beta hydroxylase of TH+ cells from control (top) and NT-1 (bottom). **P <0.01, ***P <0.001. Scale bar 2 mm and 500 µm in Fig. **a**. Scale bar 50 µm in Fig. **i, j, k** and **l**. Con – control, Con-a-control anterior, Con- m- control middle, Con-p- control-posterior, DBH – dopamine beta-hydroxylase, NM-neuromelanin pigmented neurons, NT-1-narcolepsy with cataplexy, NT-1-a: NT-1 anterior, NT-1-m: NT-1 middle, NT-1-p: NT-1 posterior. TH+ - tyrosine hydroxylase.

We then performed immunohistochemical analyses using a tyrosine hydroxylase (TH) antibody to assess norepinephrine neuronal loss (Fig. 1, g & h and i & j) (Extended Data Table. 1). A large majority of the TH containing LC neurons also had NM. Like the NM data, TH-positive cell loss in NT-1 patients was 44% (t = 8.12, df = 14, P = 0.001), and there was a significant increase in the size of TH-positive cells in NT-1 patients compared to controls (t = 7.21, df = 14, P = 0.01). The percent loss of TH+ neurons was not significantly correlated with the age of onset of cataplexy (r = 0.35, P = 0.28). Fig.1 k & l show the dopamine β-hydroxylase (DBH) stained cells in control and NT-1 patients.

We found a significant 118% increase in microglial density in the (LC) of NT-1 patients (Fig. 2a & b), (t = 4.45, df = 14, P = 0.001). Furthermore, there was a notable enlargement of LC microglia, with a 33% increase in size (t = 3.41, df = 14, P = 0.001) (Extended data Table 2). The distribution of Iba1-positive cells in both control and NT-1 subjects is shown in Figures 2c & d. Microglial clustering around neuromelanin-pigmented cells is evident in the LC of narcolepsy with cataplexy (NT-1) patients (Fig. 2e). There was no significant correlation between the increased numbers of Iba1 and the age of disease onset (r = 0.39, P = 0.24) or of disease duration (r = 0.41, P = 0.22). The significant increase in the number of microglial cells in the LC of human narcoleptics suggests a neuroinflammatory process^11^. The neuropathology report for our NT-1 patients indicated that 40% of them exhibited some degree of Alzheimer’s pathology (Extended data Table 1). Immunohistochemical study showed that most of the Hcrt axons were lost in the LC of NT-1 patients (Fig. 2 f & g), consistent with the loss of hypothalamic Hcrt neurons in narcolepsy. We investigated the presence of α-synuclein and tau protein deposits in the LC of NT-1 and control subjects. No tau or α-synuclein deposits were found in control subjects; however, tau (Fig. 2 h & i) and α-synuclein (Fig. 3 j & k) deposits were observed in the LC of NT-1 patients. Although the narcolepsy patients had differing drug treatments (Extended Data Table 1), all human narcolepsy patients showed similar brainstem pathology.

**Fig. 2:**
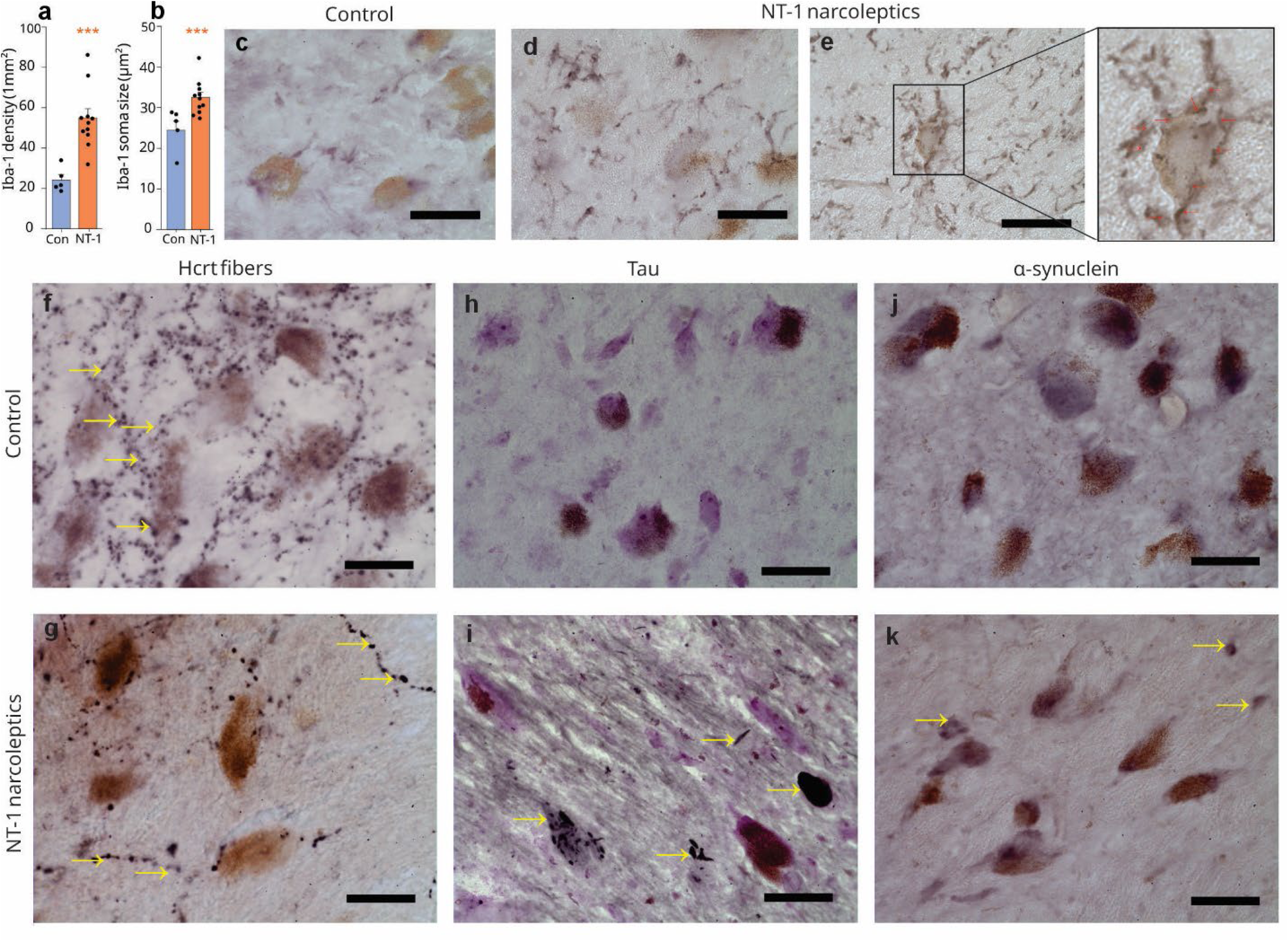
Distribution of microglia, Hcrt fibers, alpha-synuclein, and Tau proteins in the LC of NT-1. **a**, Immunohistochemistry of Iba1(ionized calcium-binding adaptor molecule) shows a significant increase in the number of microglia in the locus coeruleus of NT-1 patients compared to control (Con) subjects (n = 5 in control and 11 in NT-1 groups, t = 4.29, P = 0.001). **b**, Iba1 cell size also significantly increased (t = 3.41, P = 0.001). Data expressed as mean ± standard error. ***P <0.001. **c**, Immunobiological images of Iba1 in control and **d**, NT-1 subject. **e**, Image shows microglia (arrows) clustered around a neuromelanin pigmented neuron. Narcoleptic brains showed increased iba1 staining. **f** & **g**, Distribution of Hcrt fibers in the LC of control and NT-1 subject. As might be expected, Hcrt axons, originating in Hcrt neurons in the hypothalamus are greatly reduced in narcoleptics because of the death of hypothalamic Hcrt neurons. **h** & **i**, Immunological staining of Tau protein (black) deposits in the LC of control and NT-1 patients. **j** & **k** Images show α-synuclein distribution in the LC of control and narcoleptic subjects. α-synuclein and Tau are increased in narcoleptics. Scale bar 50 µm.

**Fig. 3:**
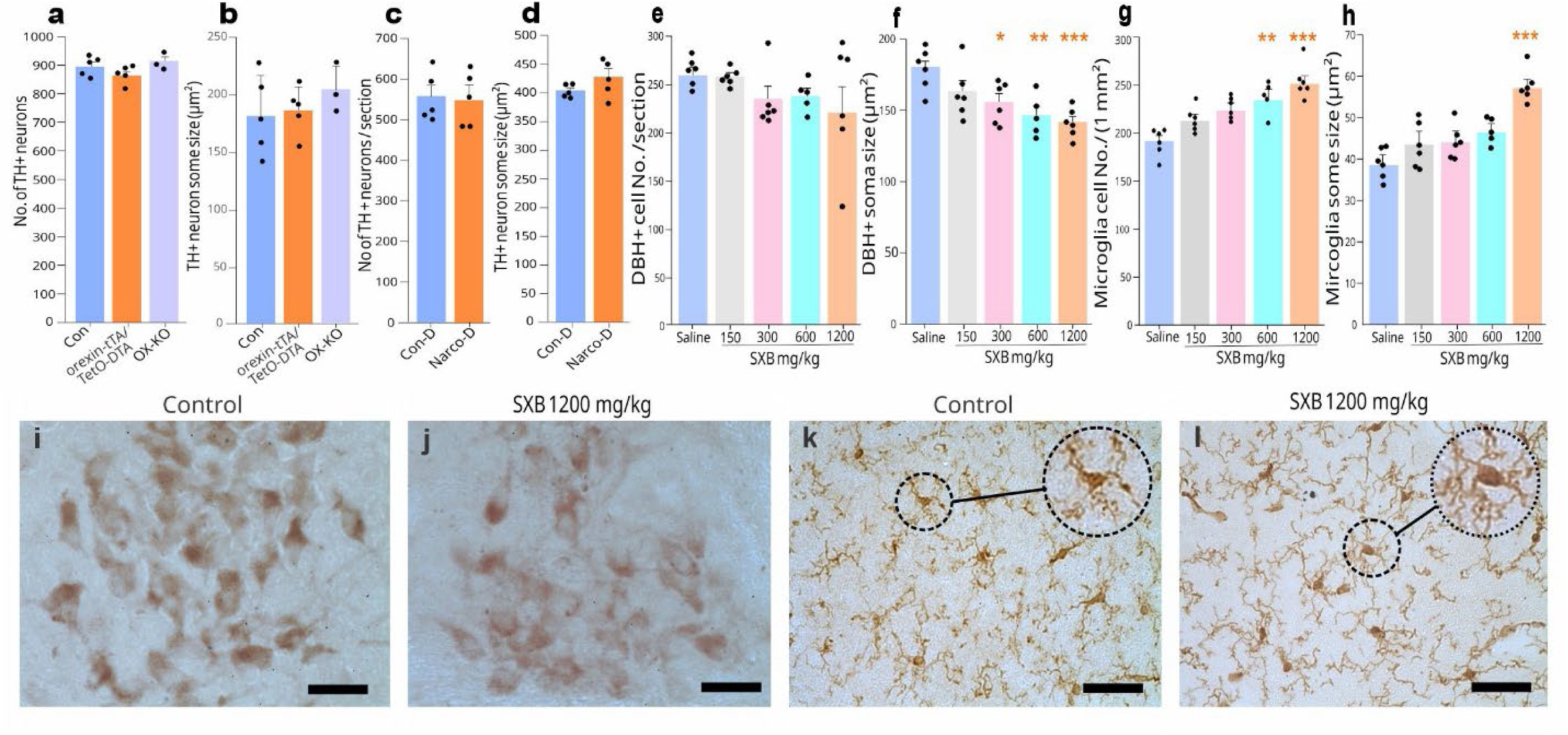
Norepinephrine neurons in mice and dog models of narcolepsy and effect of Sodium oxybate (SXB) on mice LC norepinephrine (DBH+) cells and microglia. **a**, The number of TH+ cells in the LC of control, orexin knockout mice and orexin-tTA/TetO-DTA mice. We saw no significant changes in the number of TH+ neurons in the LC in any of these models compared to controls (n = 5 in control, Orexin knockout mice N = 3 (t = 0.85, P = 0.42) and DTA group (n = 5, t = 0.17, P = 0.16). **b**, There is no difference in TH+ cell size between control and orexin knockout mice (t = 0.99, P = 0.36) or control and DTA mice (t = 0.27, P = 0.79). **c**, Narcoleptic Doberman pinschers show no difference from controls in the number of TH+ neurons in the LC (n = 5 in each group, t = 0.22, P = 0.83). **d**, TH+ cell size also did not differ between narcoleptic and control dogs (t = 1.45, P = 0.18). Data is expressed as mean ± standard error. Con-control, Con-D -control dog, LC-locus coeruleus, Narco-D-narcoleptic dog, OX-KO-orexin knockout mice, TH-tyrosine hydroxylase. **e**, In comparison to the saline group (n = 6), the number of dopamine beta hydroxylase (DBH+) cells was not significantly affected by various doses of SXB (150 mg/kg, n = 6; 300 mg/kg, n = 6; 600 mg/kg, n = 5; 1200 mg/kg, n = 6). ANOVA (df = 4, 24, F = 1.27, P = 0.31). **f**, The cell size of DBH+ decreased with higher concentrations of SXB (ANOVA, df = 4, 24, F = 6.65, P = 0.001). DBH+ cell size was significantly different from that of the saline controls at 300 mg/kg (13.7%, P = 0.047), 600 mg/kg (18.9%, P = 0.005) and 1200 mg/kg SXB (21.6%, P = 0.0001), all Tukey post-hoc. **g**, Microglia cell number (density) increased with higher doses of SXB (ANOVA, df = 4, 24, F = 7.81, P = 0.001). The increase in the number of microglia cells is significant (Tukey post-hoc) at the dose of 600 mg/kg (22.08%, P = 0.012) and 1200 mg/kg SXB (31.7%, p = 0.001). **h**, The size of microglia increased with increasing doses of SXB (ANOVA, df = 4, 24, F = 7.37, P = 0.001). A significant increase was found at 1200 mg/kg (P = 0.001, Tukey post-hoc). **i,** Histological images of DBH + neurons with saline and **j**, 1200 mg/kg SXB. **k**, Images of microglia with saline and **l**, 1200 mg/kg SXB treated mice. Data expressed as mean ± standard error. *P <0.05, **P <0.01, ***P <0.001, compared to saline group. DBH – dopamine beta-hydroxylase, SXB - sodium oxybate, Scale bar 50 µm.

We used two distinct mouse models of narcolepsy. One model was orexin-tTA/TetO-DTA (orexin-DTA) in which Hcrt neuron degeneration can be initiated by removal of doxycycline (DOX) from the diet^12^. Removing DOX food for one month caused an 87% loss of Hcrt neurons in these animals (Supplemental Fig. 1). The second mouse model of narcolepsy is mice in which the peptide cannot be synthesized^1^ (OX-KO) (Extended Data Fig. 3). We evaluated the changes in LC norepinephrine cells in these two narcoleptic models, quantifying immuno-stained TH+ cells (Fig. 3 a & b). We saw no significant changes in the number (DTA: t = 1.52, df = 8, P = 0.17, OX-KO: t = 0.85, df = 6, P = 0.42) or size (DTA: t = 0.27, df = 8, P = 0.79, OX-KO: t = 0.99, df = 6, P = 0.36) of the TH+ cells in the LC in these two animal models of narcolepsy. In addition to the two mouse models, we also examined changes in norepinephrine cells in the LC of narcoleptic dogs, a genetic narcoleptic disorder that occurs without any postnatal manipulation^13,14^ (Extended Data Table 3). No differences were observed in the norepinephrine cell number (t = 0.22, df = 8, P = 0.83) and size (t = 1.45, df = 8, P = 0.18) compared to control dogs (Fig. 3 c & d). Thus, the loss of norepinephrine in locus coeruleus that we describe here is a feature of human NT-1, not observed in any of the animal models of narcolepsy.

Sodium oxybate (SXB) is an effective treatment for human narcolepsy with cataplexy^15,16^. We conducted a dose-response study of the effect of SXB on LC in wild type mice. We used dopamine beta hydroxylase DBH to stain norepinephrine neurons and the Iba1 stain to quantify the number and size of microglial cells. The doses tested were 150 mg/kg, 300 mg/kg, 600 mg/kg, and 1200 mg/kg (Extended Data Table 3). SXB treatment had no significant effect on the number of DBH+ neurons (Fig. 3 e, ANOVA, df = 4, 24, F = 1.27, P = 0.31), but there was a dose dependent decrease in the size of DBH neurons (Fig. 3 f, ANOVA, df = 4, 24, F = 6.65, P = 0.001). Tukey post-hoc comparisons indicated a significant size reduction as compared to saline controls with 300 mg/kg (13.7%, P = 0.047), 600 mg/kg (18·9%, P = 0.005) and 1200 mg/kg SXB (21.6%, P = 0.001). Figure 3 i & j shows DBH+ neurons from saline and 1200 mg/kg SXB treated mice.

SXB significantly increased the number of microglia cells in locus coeruleus (ANOVA, df = 4, 24, F = 7.81, P = 0.001) (Fig. 3 g). The increase in the number of microglia cells was significant (Tukey post-hoc) at the dose of 600 mg/kg (22·08%, P = 0.012) and 1200 mg/kg SXB (31.7%, P = 0.001). The size of the microglia significantly increased with higher doses of SXB (ANOVA, df = 4.24, F = 7.37, P = 0.001, Fig. 3 h). A significant increase was found at 1200 mg/kg SXB (48.28%. P = 0.001). Microglia size increased in comparison to saline (Fig. 3k) with various doses of SXB, particularly at the 1200 mg/kg SXB dose (Fig. 3 l). We see LC pathological changes in all the brains of human narcoleptics we have examined, but none in the DTA^11^ or Hcrt KO mice^1^, or in the narcoleptic dogs^17^ despite the descending projection of hypothalamic Hcrt neurons to brainstem and spinal cord^18^. The findings that defined the loss of Hcrt neurons as a central feature of human narcolepsy with cataplexy, should not obscure possible differences between canine, murine and human narcolepsy. Although both murine and human narcolepsy are triggered by the loss of Hcrt neurons, in this study we find that human but not murine cataplexy is accompanied by the loss of neuromelanin containing tyrosine hydroxylase neurons in locus coeruleus. Neuromelanin is not found in rodent tyrosine hydroxylase neurons at any age^19^. The reduction of brainstem neuromelanin cells in human narcoleptics may contribute to narcoleptic symptomatology. Cells with this neurochemical composition do not exist in rodents or dogs^19^.

SXB’s therapeutic effect in humans may not be on the greatly reduced population of Hcrt neurons, but instead on norepinephrine brainstem neurons. SXB reduced TH immuno-intensity in LC^20^. Sodium oxybate has been shown to improve night-time sleep quality and reduce symptoms of both excessive daytime sleepiness and cataplexy in patients with narcolepsy. Although their precise mechanism of action is not fully understood, it is believed to primarily involve the activity of oxybate on neurons with gamma-aminobutyric acid type B (GABA-B) receptors in regions that regulate sleep-wake homeostasis^21^. Our findings indicate a dose-dependent increase in both the number and size of microglia with SXB treatment.

Tau and α-synuclein deposits in the LC of narcolepsy with cataplexy patients suggest that NT-1 shares some neurodegenerative pathology with Alzheimer’s and Parkinson’s diseases, despite earlier claims that narcolepsy decreased the incidence of Alzheimer’s^22^. However, the precise link, if any, between these findings and the mechanisms underlying narcolepsy is not clear. Important differences exist between narcolepsy and other neurodegenerative disorders. Narcolepsy symptoms typically emerge in the teenage years, in parrallel with the maturation of neuromelanin pigmentation.

In narcolepsy with cataplexy patients, there is an almost complete loss of Hcrt neurons and axons, suggesting that both Hcrt and norepinephrine contribute to the occurrence of cataplexy. We find that in the human LC, neuromelanin containing norepinephrine neurons, concentrated in the rostral locus coeruleus, are lost in narcolepsy. The rostral LC, where the loss of norepinephrine neurons is maximal, is likely to project to more rostral regions of the brain than caudal LC. One might expect that the loss of 46% of the neuromelanin containing neurons of the LC in humans with narcolepsy would produce symptoms not present in murine models of narcolepsy lacking only the Hcrt neurons of the hypothalamus, since neuromelanin neurons are not typically present in rodent brains^19^. Differences between the symptoms in these three animal models of narcolepsy and human narcolepsy would be expected. Further quantitative research, with more precise measures of motor function, and particularly muscle tone control, may clarify the behavioral consequences of the neuroanatomical changes between these differing forms of narcolepsy.

Our findings of brainstem neurodegeneration in narcolepsy with cataplexy patients demonstrates that the “Hcrt loss” explanation of human narcolepsy is incomplete. Brainstem as well as hypothalamic structures are altered in human narcolepsy. Sodium oxybate, one of the most effective treatments for narcolepsy, appears to operate on the LC region, rather than on the hypothalamus. This work identifies new targets for therapeutic interventions aimed at reducing the impact of neuroinflammation on sleep-wake regulation in narcolepsy.

## Author contributions

TCT and JMS conceptualized the study. All authors contributed to the investigation. TCT did human and animal tissue immunohistochemistry and data acquisition. MFW conducted the sodium oxybate (SXB) experiment, prepared DTA mice and HcrtR-2 mutant canines. MFW validated the statistical analysis.

Neuropathological reports were prepared by MC. JMS acquired funding and supervised the study. TCT and JMS wrote the original draft. All authors contributed to the review and editing of the manuscript.

## Data sharing

The data is available at request to the corresponding author upon publication.

## Acknowledgements

Research was supported by NIH grants DA034748, DA058639 to JMS and by the Medical Research service of the Department of Veterans Affairs to JMS. We gratefully acknowledge the brain donors and their families for making this research possible.

## Methods

### Ethics declaration

Human postmortem brain tissue was obtained under permission and used in accordance with the relevant ethical boards at VA Greater Los Angeles Healthcare System. All animal procedures were approved by the Institutional Animal Care and Use Committees at the University of California, Los Angeles, and the Veterans Administration Greater Los Angeles Healthcare System.

### Human brainstem immunohistochemical analysis

In this study, we examined post-mortem brain samples from eleven individuals with narcolepsy and five neurologically healthy controls. A summary of the brain sample characteristics is provided in Extended Data Table I. Narcolepsy diagnoses were based on the criteria set by the American Academy of Sleep Medicine: International Classification of Sleep Disorders^23^. The control group included individuals with no neurological disorder history. Brainstem tissues from narcolepsy and control subjects were obtained from the Department of Veterans Affairs Biorepository Brain Bank in Los Angeles and the Eunice Kennedy Shriver National Institute of Child Health and Human Development Brain Bank in Maryland.

Formalin-fixed brainstem tissues in one hemisphere from control and narcoleptic human patients were equilibrated in 20% and 30% sucrose solutions. Following this, 40 µm coronal sections were prepared using a freezing microtome, and sections collected at intervals of one-in-twelve. One series of sections were stained with Hematoxylin and Eosin (PK501, FD Neurotechnology Inc., Baltimore, MD) to highlight neuromelanin-pigmented cells in the locus coeruleus. Stereological techniques were employed to quantify the cell number, distribution, and size of these cells. Analysis was conducted using a Nikon E600 microscope with a three-axis motorized stage, video camera, Neurolucida interface, and Stereo Investigator software (MBF Bioscience, VT). The total number of neuromelanin-pigmented cells was then calculated for both brain hemispheres.

The neuropathological changes in the locus coeruleus (LC) were evaluated by using primary antibodies for Hcrt, TH, DBH, Iba1, Tau, and α-synuclein. Matched sections from the anterior, middle, and posterior parts of the LC were prepared based on adjacent H&E-stained sections and the Human Brainstem Reference Atlas. Free-floating sections underwent antigen retrieval before immunostaining following the method outlined by Thannickal, et al.^4, 24^. The sections were incubated in 0.5% sodium borohydride for 30 minutes. After washes with phosphate-buffered saline (0.1M, pH 7.4), transferred to 0.5% hydrogen peroxide for 30 minutes, washed, and then heated at 80°C for 30 minutes in a sodium citrate solution (10 mM, pH 8.5). The sections cooled to room temperature in the sodium citrate solution and were then washed with PBS. Following the PBS washes, the following primary antibodies were applied: Orexin A (1: 10000, rabbit polyclonal, Cat. No. H-003-30, Phoenix Pharmaceutical Inc., CA), tyrosine hydroxylase (1: 10000, rabbit polyclonal, Cat. No. AB152, Millipore Sigma, MO), dopamine beta-hydroxylase (DBH; 1:5000, rabbit monoclonal, Abcam, Cat. No. ab209487), and α-synuclein (1:5000, rabbit monoclonal, Abcam, Cat. No. ab51253). The sections were incubated in 1% normal goat serum in 0.01 M PBS for 2 hours, followed by a 72-hour incubation at 4°C with the primary antibodies. For secondary antibody, sections were incubated with biotinylated goat anti-rabbit IgG (1:500, PK-6101, Vector Laboratories) for 2 hours at room temperature, followed by avidin-biotin-peroxidase (1:300, PK-6101, Vector Laboratories) for 2 hours. The tissue-bound peroxidase was visualized using a diaminobenzidine and nickel reaction (SK-4100, Vector Laboratories). For the DBH staining, Vector SK-4600 was used to achieve a purple color. Goat polyclonal Iba1 antibody (1:10000, Abcam, Cat. No ab5076, Abcam) was used for microglia. Sections incubated with secondary antibody (1;500, (PK-6105, vector laboratories). Mouse monoclonal Tau antibody (1:5000, Cat. No. ab246808, Abcam) used for tau pathology. Sections were incubated with secondary antibodies (1:500, PK-6102, Vector laboratories) and followed by avidin-biotin-peroxidase (1:300) for 2 hours at room temperature. The tissue-bound peroxidase was visualized by diaminobenzidine and the nickel reaction (SK-4100, Vector laboratories). After staining the sections were dehydrated with alcohol, cleared with xylene and cover slipped on resinous mounting medium. The density of TH+ and Iba1+ cells was calculated as the number of cells per unit area (1 mm*²*). The “nucleator” probe in the stereology program was used to estimate the mean cross-sectional area of the cells.

## Animal models of Narcolepsy

### Immunostaining for control and mice models of narcolepsy

We conducted parallel investigations using orexin knockout (OX-KO) mice, orexin-DTA mice, and dog models of narcolepsy (Extended Data Table 3). The orexin-DTA mice were C57BL/6 mice with postnatal ablation of Hcrt neurons, which was induced by doxycycline withdrawal. The experimental group had doxycycline (DOX) food removed for 30 days, followed by a reintroduction of DOX food and euthanasia 14 days later. Control animals were administered DOX food throughout and were sacrificed at the same time as the experimental group. Control and both narcolepsy mouse models were perfused, post-fixed, and transferred through 20% sucrose and 30% sucrose solutions. No antigen retrieval was necessary for the mouse sections. Brainstem sections were cut at 40 µm, and a one-in-three series was analyzed. The sections were blocked in 1% normal goat serum in 0.01 M PBS for 2 hours and then incubated with tyrosine hydroxylase (1:10,000, rabbit polyclonal, Cat. No. AB 152, Millipore Sigma, MO) for 72 hours at 4°C. Sections were subsequently incubated with a secondary antibody (1:500, biotinylated goat anti-rabbit IgG, PK-6101, Vector Laboratories) followed by avidin-biotin-peroxidase (1:300, PK-6101, Vector Laboratories) for 2 hours each at room temperature. Peroxidase activity was visualized with a diaminobenzidine reaction (SK-4100, Vector Laboratories). Tyrosine hydroxylase-positive (TH+) neurons were bilaterally counted in a one-in-three series, and the count was reported without multiplication by 3. Stereological analysis using a “nucleator” probe was performed to assess cell number, distribution, and size, using a Nikon E600 microscope equipped with a three-axis motorized stage, video camera, Neurolucida interface, and Stereo Investigator software (MBF Bioscience, VT).

### Immunostaining for control and narcoleptic dogs

Narcoleptic (Hcrt-R2 mutant) Doberman pinschers and breed-matched controls were studied (Extended Data Table 3). The animals were perfused with saline and formalin, and their brains were removed and stored in formalin prior to sectioning and staining. The formalin-fixed brainstem was infiltrated with 20% and 30% sucrose. Coronal sections (40 µm) were cut on a freezing microtome with one-in-twelve section intervals. One series of sections was stained with crystal violet (PK501, FD Neurotechnology Inc., Baltimore, MD). The sodium citrate heat antigen retrieval method was used before doing TH immunostaining. For tyrosine hydroxylase (TH) immunostaining, data acquisition and analysis followed the same protocol used for the mice. TH+ cells were counted bilaterally, and the data was reported as the number of neurons per section.

### Effect of Sodium Oxybate on Norepinephrine and Microglia Cells in Mice

The animals used in this study were part of the research published by Wu et al^20^. Mice were treated with either saline or sodium oxybate (sodium oxybate oral solution, 500 mg/mL, Xyrem, Jazz Pharmaceuticals) at doses of 150, 300, 600, or 1200 mg/kg, administered intraperitoneally (IP) for 14 days (Extended Data Table 3). Injections at each dose were given twice daily, at ZT 2 (09:00 am) and ZT 6 (01:00 pm). Two hours after the final dose on the 14th day of treatment, the animals were euthanized with Fatal Plus (100 mg/kg), followed by transcardial perfusion with saline and then 4% paraformaldehyde. The brains were carefully removed from the skull and immersed in 4% paraformaldehyde for 24 hours. After fixation, the brains were transferred to 20% sucrose until they sank, then placed 30% sucrose for 3 days. The brainstems were sectioned into 40 μm slices using a freezing microtome. The sections were collected through one-in-three section intervals. For immunohistochemistry, brain sections were washed in PBS and treated with 0.5% hydrogen peroxide for 30 minutes, followed by PBS washes. Sections were then incubated in blocking solution (1% normal goat serum and 0.3% TritonX-100 in PBS) for 2 hours and subsequently incubated with dopamine beta-hydroxylase antibody (1:5000, rabbit monoclonal, Abcam, Cat. No. ab209487) for 72 hours at 4°C. Afterward, sections were incubated with biotinylated goat anti-rabbit IgG (1:500, PK-6101, Vector Laboratories) for 2 hours at room temperature, followed by avidin-biotin-peroxidase (1:300, PK-6101, Vector Laboratories) for 2 hours. The tissue-bound peroxidase activity was visualized using a diaminobenzidine reaction (SK-4100, Vector Laboratories).

Data analyses were performed by blind to treatment conditions. All staining, counting, and cell measurements were done on coded tissue. In every case, the same individual counted experimental and control tissues in randomized order. Control sections from each brain were processed without the primary antibody and did not show staining. Human brainstem regions and nuclei were identified using the “Atlas of the Human Brain”^25^. For mice, the “Mouse Brain in Stereotaxic Coordinates”^26^, and for dogs, the “Stereotaxic Atlas of The Dog’s Brain”^27^. Digital images were captured using a Micro Fire camera (Optronics, Goleta, CA) and imported into Corel Draw for contrast and brightness adjustments were made as needed.

### Statistical analysis

All statistical analyses were performed by SYSTAT Version 13. Data are presented as mean ± S.E.M. The number of subjects in each human and animal study is listed in Extended Table 1, as well as in Extended Table 2 and Extended Table 3. A one-way ANOVA was used to assess the dose-response of SXB, followed by Tukey post-hoc comparisons among different doses. Pearson’s correlation coefficient (r) was used to examine the relationship between cell loss, disease onset, and duration. Two-group comparisons were made using the t-test.

## Additional information - Extended Data

**Extended Data Table 1:**
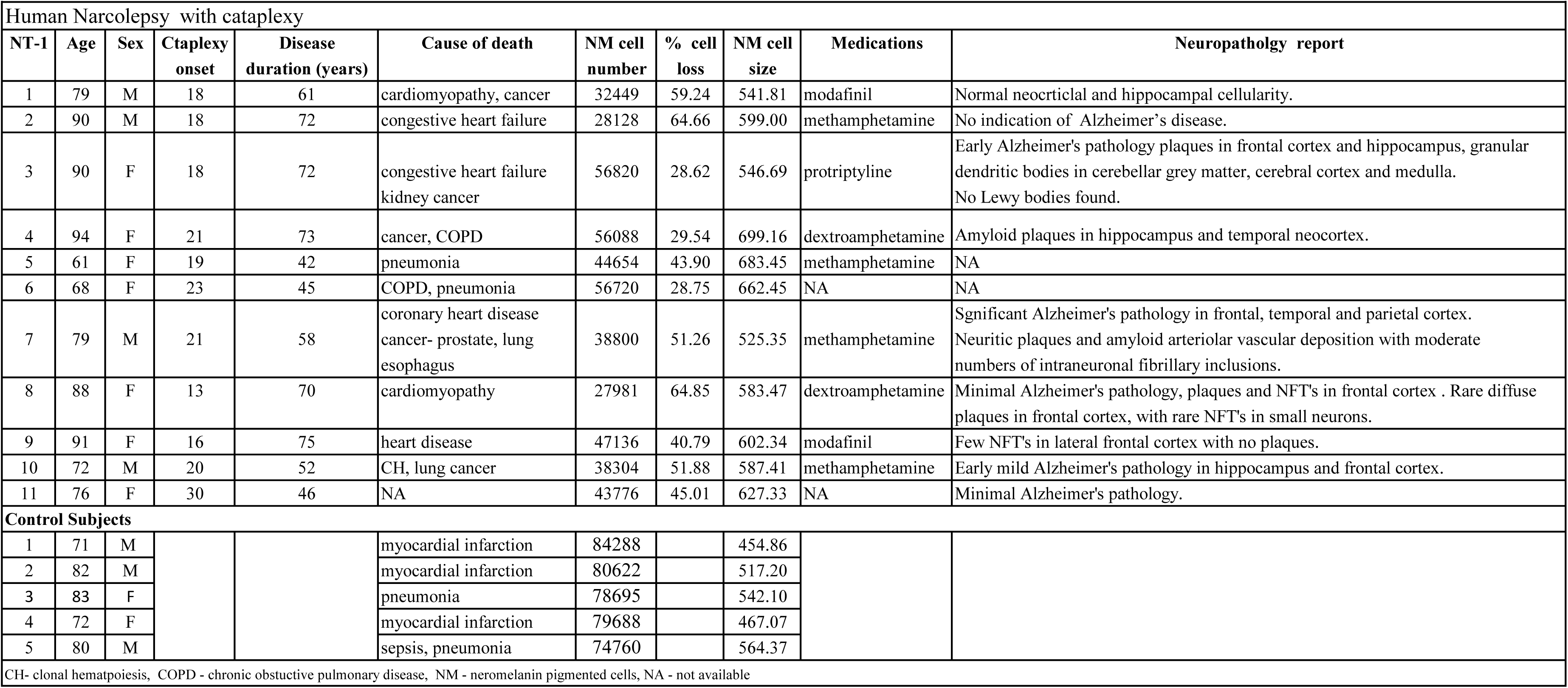
The characteristics of Human subjects and neuromelanin pigmented cell number and size.

**Extended Data Table 2:**
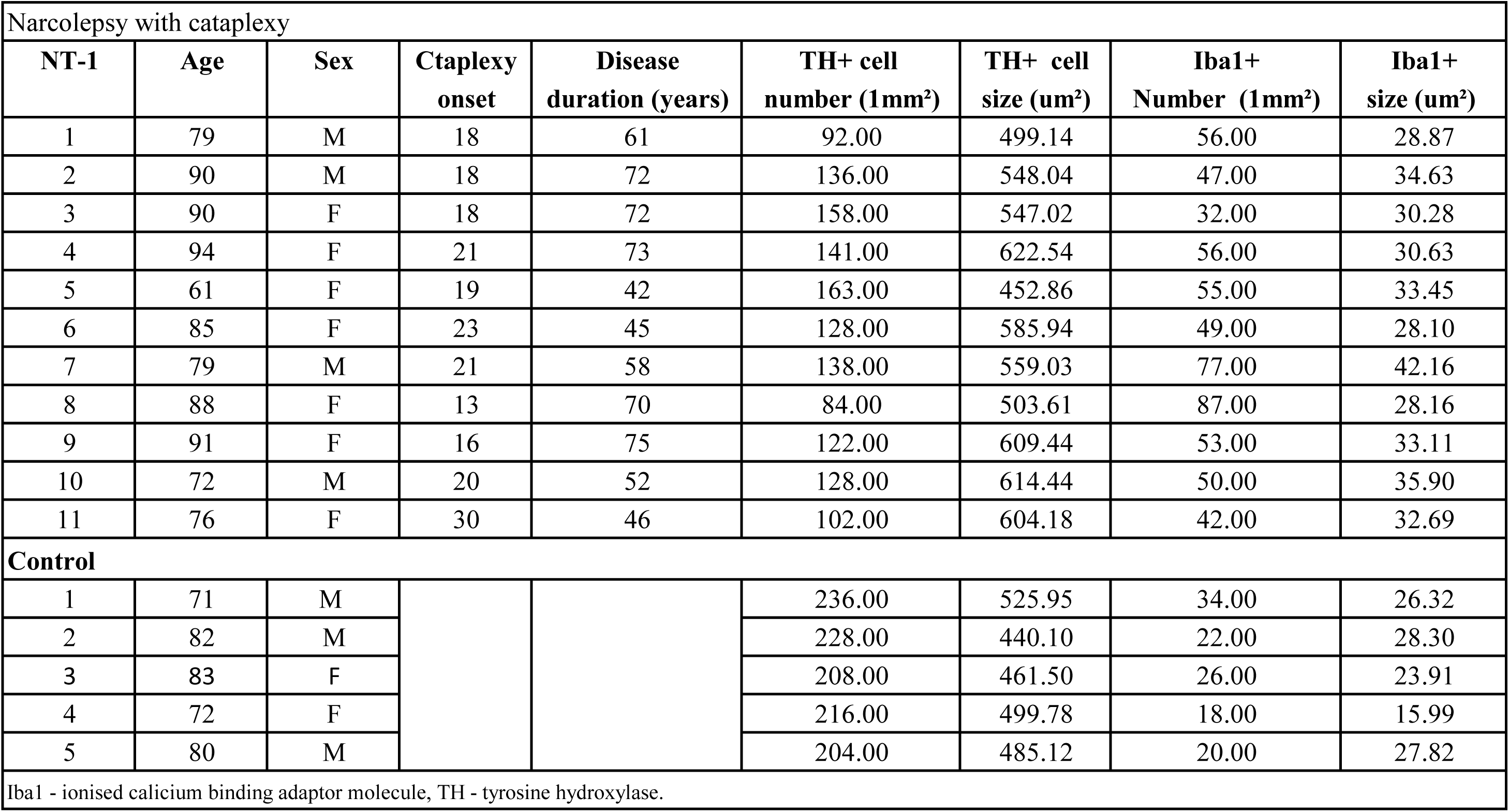
The number and size of TH and Iba1 cells in human narcolepsy and control subjects.

|  |
| --- |
| Narcolepsy with cataplexy |

| NT-1 | Age | Sex | Ctaplexy onset | Disease duration (years) | TH+ cell number (1mm<sup>2</sup>) | TH+ cell size (um<sup>2</sup>) | Iba1+ Number (1mm<sup>2</sup>) | Iba1+ size (um<sup>2</sup>) |
| --- | --- | --- | --- | --- | --- | --- | --- | --- |
| 1 | 79 | M | 18 | 61 | 92.00 | 499.14 | 56.00 | 28.87 |
| 2 | 90 | M | 18 | 72 | 136.00 | 548.04 | 47.00 | 34.63 |
| 3 | 90 | F | 18 | 72 | 158.00 | 547.02 | 32.00 | 30.28 |
| 4 | 94 | F | 21 | 73 | 141.00 | 622.54 | 56.00 | 30.63 |
| 5 | 61 | F | 19 | 42 | 163.00 | 452.86 | 55.00 | 33.45 |
| 6 | 85 | F | 23 | 45 | 128.00 | 585.94 | 49.00 | 28.10 |
| 7 | 79 | M | 21 | 58 | 138.00 | 559.03 | 77.00 | 42.16 |
| 8 | 88 | F | 13 | 70 | 84.00 | 503.61 | 87.00 | 28.16 |
| 9 | 91 | F | 16 | 75 | 122.00 | 609.44 | 53.00 | 33.11 |
| 10 | 72 | M | 20 | 52 | 128.00 | 614.44 | 50.00 | 35.90 |
| 11 | 76 | F | 30 | 46 | 102.00 | 604.18 | 42.00 | 32.69 |

Control
|  |  |  |  |  |  |  |  |  |
| --- | --- | --- | --- | --- | --- | --- | --- | --- |
| 1 | 71 | M |  |  | 236.00 | 525.95 | 34.00 | 26.32 |
| 2 | 82 | M |  |  | 228.00 | 440.10 | 22.00 | 28.30 |
| 3 | 83 | F |  |  | 208.00 | 461.50 | 26.00 | 23.91 |
| 4 | 72 | F |  |  | 216.00 | 499.78 | 18.00 | 15.99 |
| 5 | 80 | M |  |  | 204.00 | 485.12 | 20.00 | 27.82 |

Iba1 - ionised calcium binding adaptor molecule, TH - tyrosine hydroxylase.

**Extended Data Table 3:**
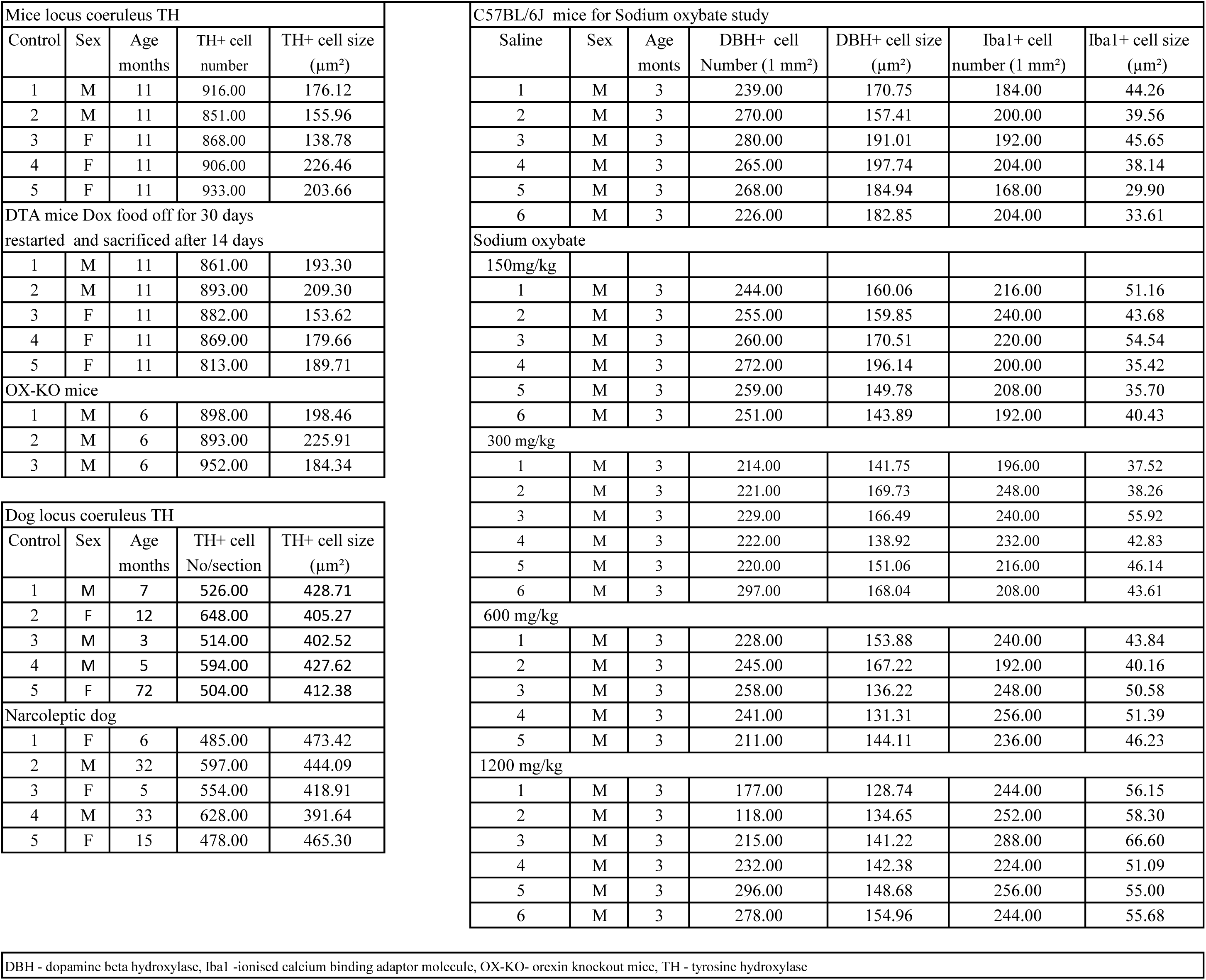
The number and size of TH+ cells in animal models of narcolepsy and effect of SXB on LC norepinephrine and microglia cells.

