## Supplemental Information figure for "Brainstem pathology in human narcolepsy: Neurodegeneration in the locus coeruleus in narcoleptic humans, but not in genetically narcoleptic mice or dogs"

Supplementary information

Supplemetary Fig.1 Hcrt cell loss in DTA mice with 30 days removal of DOX food.

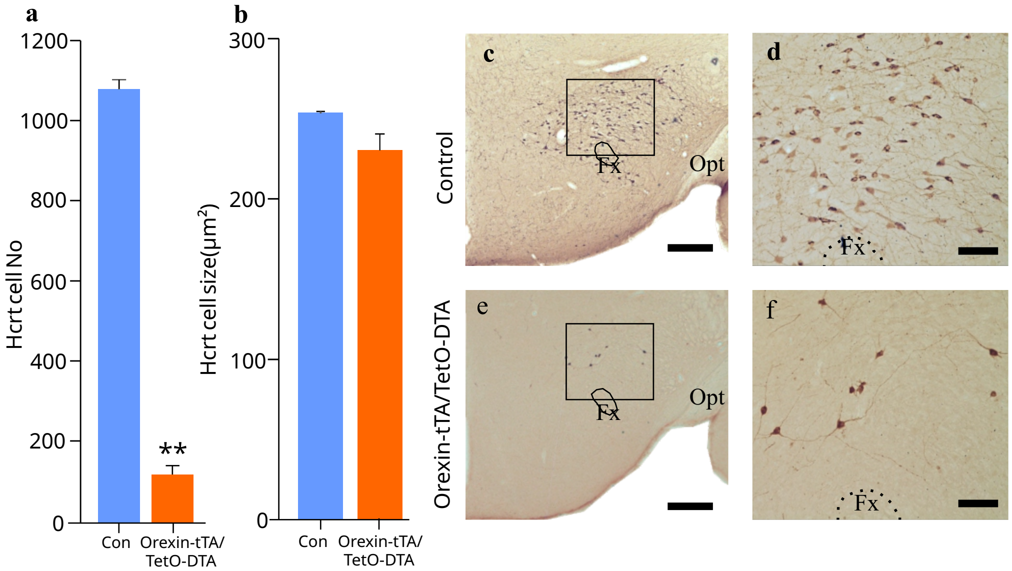


Fig. S1. **a**, 90% Hcrt cell loss in DTA mice (control N = 2, experimental N = 5, t = 25.5, df = 5, P = 0.01). **b**, There is no difference in cell size (t = 1.4, df = 5, P = 0.22). **c** & **d**, Histological images from control. **e** & **f**.,Images from DTA mice with hypocretin loss. Scale bar 500 μm (**c** & **e**) and 100 μm (**d** & **f**).
